# A GWAS platform built on iPlant cyber-infrastructure

**DOI:** 10.1101/002881

**Authors:** Liya Wang, Doreen Ware, Carol Lushbough, Nirav Merchant, Lincoln Stein

## Abstract

We demonstrated a flexible Genome-Wide Association Study (GWAS) platform built upon the iPlant Collaborative Cyber-infrastructure. The platform supports big data management, sharing, and large scale study of both genotype and phenotype data on clusters. End users can add their own analysis tools, and create customized analysis workflows through the graphical user interfaces in both iPlant Discovery Environment and BioExtract server.

## I. INTRODUCTION

The iPlant Collaborative (iPlant) is a United States National Science Foundation (NSF) funded project that has created an innovative, comprehensive, and foundational cyber-infrastructure (CI) in support of plant biology research [1]. iPlant is an open-source project with application programming interfaces that allow the community to extend the infrastructure to meet their needs.

A Genome-Wide Association (GWA) Study is an examination of many common genetic variants in different individuals to see if any variant (e.g. single-nucleotide polymorphisms (SNPs), insertion-deletions (indels), etc.) is associated with a trait (e.g. disease, plant height, etc.) [2]. A typical GWA study will require genotyping a group of individuals, variant detection and imputation, phenotyping the same group, trait data extraction, and association mapping. With rapid advances in genotyping and phenotyping techniques in the last decade, large amounts of data are accumulating. The storage, sharing, analysis, and re-analysis of these data have become a challenge in biology research. Therefore, GWA is closely tied to the mission of iPlant for building a research oriented CI, especially in modeling complicated data types for storage, sharing and analysis.

Here, we present a cloud-based open platform built on iPlant CI for GWA. The platform supports parallel computation on high performance computing clusters (cloud-based) and open for end-users to add new analysis tools to extend the analysis. The analysis tools and workflows presented are accessible through the graphical user interfaces in both the iPlant Discovery Environment and the BioExtract server [3].

## II. BACKGROUND

Conventional linkage or quantitative trait loci (QTL) mapping has been effective in identifying genes associated with traits of interest. However, the identified genes are commonly restricted to the ones segregating in the Recombinant Inbred Line (RIL) families. Additionally, the mapping resolution is usually low due to the limited number of recombination events that occur during the creation of the RILs. On the other hand, in most cases, a GWA study is performed with natural populations by scanning an entire genome for SNPs associated with a trait of interest, thus enabling mapping quantitative traits with high resolution in a way that is statistically very powerful.

A major issue of GWA mapping is that it generates false positives due to population structure [4], a systematic difference in allele frequencies between subpopulations in a population possibly due to different ancestry. One commonly used approach for controlling population structure is structural association (SA), which relies on randomly selected markers (or SNPs) from the genome to estimate population structure. The estimated population structure, e.g. using STRUCTURE [5], is then incorporated into the association analysis. To further capture the relatedness between individuals for eliminating more false positives, the unified mixed model approach has been proposed, including different MLM (mixed linear model) approaches implemented in TASSEL [6, 7] and EMMAX [8]:

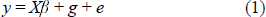

where *y* is the phenotype, *X* is a matrix of fixed effects, including genotypes and population structure, *β* is a vector of effect size, *g* represents random effects, including the matrix of kinship coefficients and the vector of polygenic effects and *e* is a vector of random independent effects with variance modeling the residual error. Both *g* and *e* are assumed to have a Gaussian distribution with a mean of zero. The general linear model (GLM), discussed later, models the associations in a fashion similar to equation (1) but without *g*.

Both TASSEL and EMMAX are based on single-locus tests combined with some kind of diffuse genomic background. For complex traits controlled by several large effect loci, these single-locus approaches may not be appropriate. Explicit use of multiple cofactors has been shown to outperform simple interval mapping in traditional linkage mapping, including both multiple-QTL mapping [9] and composite interval mapping, e.g. QTL Cartographer [10]. Therefore, a multi-locus mixed model approach, MLMM, has been proposed with a stepwise regression strategy and shown to be more sensitive (fewer false negatives) than single-locus approaches on both simulation and real data set [11]. Therefore, during the course of building the iPlant CI, we have integrated QTL Cartographer, STRUCTURE, TASSEL, EMMAX, MLMM, and many other applications for adding key functionalities supporting association analysis. We also developed in-house wrapper scripts for connecting these applications through various format conversions, thus enabling an automated reusable workflow for large scale GWA studies.

iPlant’s scope is beyond GWA but closely tied to GWA. A large portion of the analysis tools integrated into iPlant CI are focused on the next generation sequencing (NGS) data analysis, including transcriptome data analysis (e.g. RNA-seq, assembly), genome assembly, epigenome data analysis (e.g. Methyl-seq, Chip-seq), and genome assembly and annotation. These developments are all related to GWA either directly or indirectly, e.g. variant calling from transcriptome data. Sequencing has become the standard method for acquiring genotypes for GWA. However, for the purpose of GWA study, at least hundreds of lines (individuals) need to be sequenced and analyzed. To reduce the sequencing cost, a simplified Genotyping-By-Sequencing (GBS) approach has been developed [12], which relies on the selection of methylation-sensitive or insensitive restriction enzymes to avoid sequencing repetitive regions. GBS significantly reduces the cost of GWA genotyping with the tradeoff of a higher degree of missing SNPs compared to whole genome sequencing. Therefore, we have integrated both the GBS analysis workflow [12] and a missing SNP imputation tool, called NPUTE [13], to support the genotype analysis for GWA.

## III. TECHNICAL DETAILS

Before starting a scientific data analysis, users need to upload the input data into iPlant Data Store, which is built on the Rule-Oritented Data System (iRODS) [14]. The most common way to upload the data is through the iPlant Discovery Environment (DE) [15]. The iPlant DE also provides integrated scientific tools that can retrieve the input data, and send the computation task to the high performance computing clusters at Texas Advanced Computing Center (TACC) via the iPlant Foundation API (fAPI) [16]. Alternatively, users can also access the input data and perform the analysis through either the iPlant Atmosphere (a virtual machine platform) or the BioExtract server [3] (alternative GUI similar to the iPlant DE). More technical details are provided in following sections.

### A. Data Store

The iPlant Data Store is a centralized facility to address the existing needs of the community to share and store scientifically relevant data sets and metadata. Underlying the Data Store is a federated network of Integrated Rule-Oriented Data System (iRODS) [14] servers running at University of Arizona and replicated at the TACC. Users can access the Data Store in several ways, including the iPlant DE [15], a RESTful web service interface through the iPlant fAPI [16], Davis web application [17] (web interfaces), iDrop [18] (desktop application), and FUSE interface [19] for command line tools. The provenance in the Data Store is addressed through the use of universally unique identifiers (UUID) for every file, folder, and piece of metadata. Every action taken by a user is associated with one or more UUID and logged by a centralized tracking service.

The instructions for registering for an iPlant account, deploying applications, and the tutorial for this workflow can be found on the public wiki site: https://pods.iplantc.org/wiki/. Once registered, the user gets a 100 GB (gigabytes) initial allocation in the iPlant Data Store, which can be increased upon request. Users can share data with collaborators through the iPlant DE and web links. The easiest way to get familiar with the iPlant CI (Data, Apps, and Analysis) is through the DE (https://de.iplantc.org/de). User support is available through.

### B. Foundation API

The iPlant fAPI is a hosted, Software-as-a-Service (SaaS) resource for the computational biology field. Operating as a set of RESTful web services, the fAPI bridges the gap between the HPC and web worlds that allows modern applications to interact with the underlying infrastructure [16]. To deploy a new software package as a private tool through fAPI that runs on the command line, a user simply needs to upload it to iPlant Data Store, and register it with a Javascript Object Notation (JSON) file through fAPI. The JSON file contains metadata describing the Graphical user interface (GUI) and the computing environment. The GUI makes entering or adjusting parameters easy for analysis or re-analysis. At runtime, the data is copied from iPlant Data Store to cluster nodes and executed with designated binaries and parameters. Results are copied back to the Data Store once completed. The private tools can be made public by notifying iPlant staffs for sharing with others.

The fAPI has been adopted by iPlant’s own DE, the BioExtract Server, and Easy Terminal Alternative [20].

### C. Discovery Environment

The iPlant DE is a distributed, service-oriented architecture (SOA) that exposes service endpoints in a RESTful manner and primarily communicates using JSON. The focus of the DE centers on providing a “software workbench” or “platform” for the execution and management of scientific analyses. GUIs for scientific analysis are driven by metadata “descriptions” encoded in a different JSON file other than the fAPI. These “descriptions” can be authored by users through a tool integration service (TiTo), enabling the collaborative extension of the overall application’s analysis capabilities. The execution of an analysis can be tailored to the needs of the computation and can either run on a local Condor cluster or a remote computing resource through the fAPI Data, Authentication (Auth), Applications (Apps), and Jobs services.

Currently, there are over 400 public applications integrated into the iPlant DE. Users can deploy their own applications by either requesting installation on the iPlant server or through fAPI by following instructions on the wiki site. Using both public and private apps, users can construct automated workflows in the DE with the interface shown in Fig. 1. This shows the three steps for constructing the variant calling workflow using the Genotyping-By-Sequencing protocol [12]: describe, add and order apps, and map outputs/inputs among apps. This workflow takes short read sequencing data and outputs variants for GWAS. The construction procedure requires that inputs and outputs of each step be explicitly named, giving the advantage that the resulting workflow can be built beforehand and reused. Once completed the workflow can be kept private or submitted for public use.

**Fig. 1.**
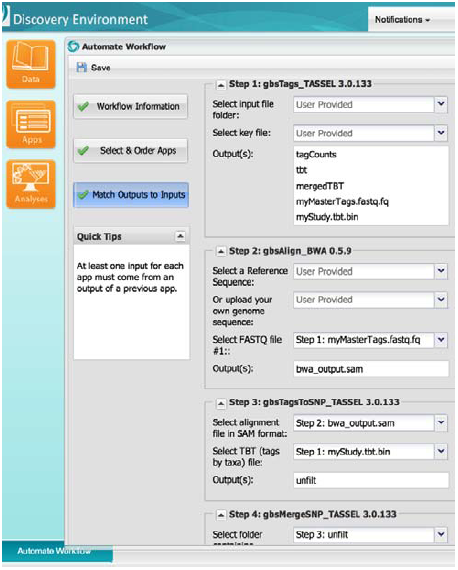
iPlant DE interface for constructing automated workflow.

### D. BioExtract

The BioExtract Server [3] is an open, web-based system designed to aid researchers in the analysis of genomic data by providing a platform to facilitate the creation of bioinformatics workflows. There is a unified authentication mechanism between iPlant and BioExtract using the fAPI’s Auth service. Therefore, any analysis tools registered through the iPlant fAPI can be accessed in BioExtract via an auto-generated GUI defined by the fAPI JSON file. Different from the DE’s build-before-run workflow construction, BioExtract implemented a run-then-build approach. That is, BioExtract’s scientific workflows are created by recording tasks performed by the user. These tasks may include execution database queries, saving query results as searchable data extracts, and executing local and web-accessible analytic tools. The series of recorded tasks can then be saved as a reproducible, sharable workflow available for future execution with the original or modified inputs and parameters.

Data in the iPlant Data Store can be accessed by BioExtract applications or workflows directly through the integrated iPlant fAPI IO and Data services. The analysis runs across systems at TACC using the iPlant fAPI job services. The public fAPI based apps of iPlant are also available on BioExtract server (under Tools/iPlant at http://bioextract.org). To use these apps and access the iPlant Data Store, user’s iPlant credentials need to be synchronized on the BioExtract server by “Register iPlant Account” (http://bioextract.org/users/create-account.xhtml). Once synchronized, all of iPlant’s public as well as the users own private apps can be used to create automated workflows. After logging in, users can click on the “Workflow” button on the top bar, then “Create and Import Workflows” and “Record Workflow”. All subsequent analyses will be recorded and can be saved as a workflow for future use.

Fig. 2 shows the execution of the GWA mapping workflow, which submits 29 jobs to TACC’s Lonestar cluster with one click and returns analysis results from six different association models. Further details are discussed in the next section. Each rectangular button on the chart represents one stand-alone tool corresponding to names in the left panel. The workflow is constructed by running each tool at least once. The connection between tools is established by using outputs of one tool as inputs for another tool. The same analysis can be repeated with different arguments in the construction process, and the failed or inappropriate ones can be deleted later. Finally,, the workflow can be shared and re-executed with different configurations of input data and parameters.

**Fig. 2.**
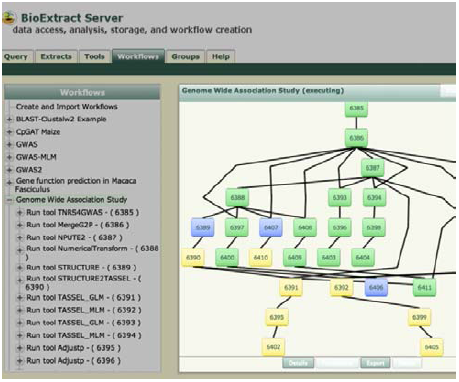
Screen shot of executing a GWAS workflow on the BioExtract server. The status of each application (icon) is distinguished by its color (green: completed; blue: executing; yellow: waiting).

### E. Atmosphere

iPlant Atmosphere provides the computing resource in a virtual machine (VM). The completely customizable VM can be launched and managed through a rich web interface. Thus, researchers can use the VM to develop novel analytical tools and provide the tools for accessing through a web browser. The VM can also be mounted to the iPlant Data Store so that the user can access their data directly for post-analysis, if necessary, after being processed in the iPlant DE.

As an example of using Atmosphere to provide GWA phenotyping tools for community use, a virtual image is built with a pre-installed application, HypoTrace [21], which is an image analysis based phenotyping tool. Such applications usually need a built-in graphical user interface for interactive operations on raw image data. The computationally demanding non-interactive operations can be passed to high performance computing clusters via the iPlant fAPI within the Atmosphere image. Users can initiate his/her own VM from the web interface using the image instance HypoTrace for analysis.

## IV. GENERAL GWA WORKFLOW

The existing GWA platforms, e.g., easyGWAS [22], focus on association mapping between prepared variant data and trait data, using predefined association tools and workflows. The iPlant CI based platform has the following key advantages. First, it provides big data management and sharing. Second, it provides large scale computational support for variant calling, imputation, and association analysis. Third, it is open for the community to contribute new analysis tools. Fourth, it supports many popular applications and provides automated workflows for GWA mapping with different models. For example, Fig. 3 shows iPlant’s general workflow for GWA. Users can construct their own workflow to suit their research needs or use their favorite association applications. The workflow presented in Fig. 3 is a mega-workflow consisting of several workflows, including the variant calling workflow (e.g. GBS workflow as shown in Fig. 1), association workflow (e.g. MLM workflow in Fig. 2), and phenotyping analysis workflow (e.g. using HypoTrace Atmosphere image).

**Fig. 3.**
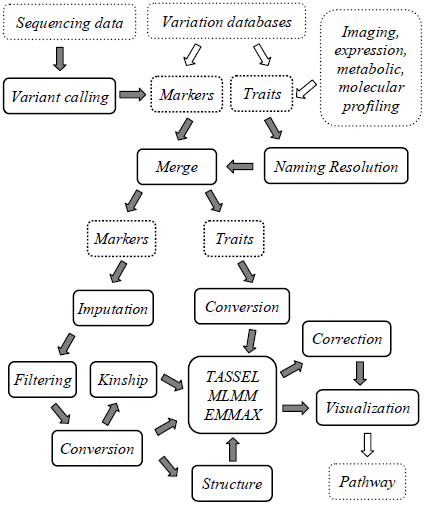
General GWAS workflow (data in a dashed box; applications in a solid box; open arrow for functionality to be built; close arrow for implemented functionality).

We will use the Sorghum Association Panel (SAP) re-sequencing data [23] as an example to walk through the workflow. The sequencing data is deposited into the NCBI SRA with accession numbers SRR636574 and SRR636575. The workflow starts in the iPlant DE by importing FASTQ sequence data into the Data Store [24] using the “NCBI SRA import” and “NCBI SRA Toolkit fastq-dump” tools. As described before, each file or folder will be assigned a UUID. The FASTQ files are further organized for feeding into the GBS workflow (built in DE) for calling variants. In this case, trait data are extracted from a previous study labeled with accession names [25]. Before merging trait data with variant data, accession names in trait data are converted to accession ids with an application named “TNRS4GWAS” for naming resolution. When users launch the app TNRS4GWAS, the users credentials are validated via the fAPI Auth service, then the input data are copied to the TACC computing clusters via the iRODS (the fAPI Data service), The fAPI Job service is then activated to submit the job to the clusters with arguments defined by TNRS4GWAS GUI through the fAPI App service.

It is essential to impute missing marker data before the association mapping. An imputation tool, NPUTE [13], is used to fill missing marker data using the nearest neighbor algorithm. Then the marker data is filtered by minor allele frequency and converted for downstream analysis. To deal with the confounding factor of population structure in GWAS, a popular application, STRUCTURE [5], is integrated. Similarly, various kinship estimation methods are added for the similar purpose of feeding into various association models.

For association mapping, we have streamlined three popular tools, including TASSEL [6–7], EMMAX [8], and MLMM [11] for comparison. Each of these tools takes marker and trait data in different formats; therefore conversion is usually needed for automating the analysis workflow. For TASSEL, different models can be tested by including population structure, kinship or not. The association between marker and trait is quantified with p values. The p values need to be corrected or adjusted for multiple comparisons. A XYPlot function is added to plot the so-called “Manhattan Plot” for visualization. Further development of the visualization function will allow interactive selection of significant p values for extracting nearby genes and their associated pathways for further analysis. For high throughput phenotyping, image analysis tool, HYPOTrace [21], has been added and will be scaled to support extracting traits from images.

NPUTE determines the optimal window size for imputation by testing known markers with various sizes of windows. Similarly, STRUCTURE picks the optimal number of clusters by testing various assumptions of the number of population clusters. These computations are time consuming but without data dependency among them. Thus, both NPUTE and STRUCTURE are integrated with the fAPI and parallelized through TACC’s parametric launcher module.

All applications are integrated in the DE for running on either a Condor cluster (Variant calling workflow) or the TACC Lonestar cluster (all GWAS applications) through fAPI. A GWAS workflow is also automated on the BioExtract Server, which allows testing of various models in a parallel fashion once the data dependency is clear. The construction of automated workflows on the BioExtract Server using fAPI based apps demonstrates that the iPlant CI makes building innovative solutions with ease, without having to worry about the foundational infrastructure.

## V. USE CASES

No single workflow can address all research problems, but our goal is to demonstrate that the iPlant platform is flexible enough for helping scientists solve real world problems. For this reason, we have two uses cases below to show how different challenges are addressed using the iPlant infrastructure. The first one is the re-analysis of the published sorghum association panel or SAP data set with the workflow presented in Fig. 2 and Fig. 3. The second one is the large scale analysis of the maize Nested Association Mapping (NAM) data set [26] with 115 traits using the bootstrapping method implemented in TASSEL. The main challenges in the first use case include extracting large amounts of sequencing data from the NCBI SRA, variant calling, imputation and automating the analysis workflow for testing various association models. The second use case involves a computationally demanding problem, bootstrapping. The entire analysis is completed in a few weeks using iPlant CI while it is estimated to require 2.5 years on a typical 8-core desktop.

Another reason we include the second use case is that the NAM data set relies on projecting SNPs from founder lines at run time, instead of keeping all SNPs beforehand, for efficient storage and association computation. It is worthwhile to point out that the iPlant platform is not designed to be a data repository of published or authoritative data, but rather, a scratch space where raw or published data can be imported and analyzed. Therefore, in the first use case, we aim to demonstrate how we extract a published data set from a public data repository, format it correctly, and combine the data with trait data for association analysis. On the contrary, the second use case shows that some popular data sets, e.g. the NAM genotype data set are hosted in the iPlant Data Store. In such cases, the user only needs to provide trait data for association mapping.

### A. Sorghum Association Panel (SAP)

Sorghum is an important crop species for feeding 500 million people in sub-Saharan Africa and south Asia because it can sustain high yields where precipitation is low or erratic. To examine the diversity of sorghum, genetically dissect its agro-climatic traits, and enable marker-assisted breeding, 971 sorghum accessions have been recently sequenced [23] using the GBS protocol [12]. After filtering for local linkage disequilibrium and tag coverage (a GBS workflow parameter describing sequencing depth), this study yielded 265,487 SNPs with an average density of one SNP per 2.7 kbp. The authors deposited around 200 GB of compressed raw sequence data (generated on the Illumina HiSeq 1000) into NCBI SRA after publication. Once uncompressed, the whole data set reaches over 1 TB (terabytes) that can be stored and analyzed using iPlant CI. The SAP data (336 accessions) is buried inside this data set.

For the GBS sequencing, barcoded multiple plant accessions were pooled together and sequenced. A single text file, called key file, contains the mapping information between barcodes and samples, including Illumina flowcell name, lane, barcode, and DNASample (accession id) on each row that are needed by the downstream GBS workflow for variant calling. The GBS workflow takes each lane as an input file and requires flowcell name (presented with instrument name instead in these deposited reads) and lane number being encoded in the file name for de-multiplexing at run time. However, NCBI requires the raw data being de-multiplexed before submission for treating each strain as a BioSample. The automatic de-multiplexing (in this case) also completely messed up the read data, making it almost impossible to repeat the variant calling analysis with the GBS workflow. After personal communication with the authors, we were able to extract the SAP data and format them properly by mapping instrument name with flowcell names. Currently, NCBI made an exception on GBS data that allows treating each lane as a BioSample. The key file should also be posted as comments that can be retrieved as the page source from the SRA website. The GBS data will be serving as a good user case for iPlant’s future developments on the data management system.

Before dealing with the missing SNPs (mostly due to the reduced representation of the genome using the GBS protocol), there is a practical problem that needs to be addressed for physically linking the genotype data [23] to available traits [25]. For the SAP data set, the SNPs are marked with the accession ids (unique identifier for each individual plant) while non-matching alias names were assigned to each accession for the trait data collected years ago. To make matters worse some plant accessions have multiple alias names used by different research groups that conducted trait research on the sorghum panel, mostly due to lacking standard naming conventions. This has created a serious problem in data integration, or in this case, linking genotype data to phenotype data. Similar to the iPlant TNRS [27] effort, we have manually curated a database for mapping various forms of accession names to unique accession ids, and developed a tool, TNRS4GWAS, to facilitate the integration of genotype data and phenotype data. Currently, the tool supports accession name to id conversion for various plant species, including Arabidopsis, Rice, Sorghum, and Maize.

As discussed earlier, we integrated a parallel version of NPUTE for imputing missing SNPs. Fig. 4A shows the imputation accuracy at different window sizes for the nearest neighbor estimation. We parallelized NPUTE so that the imputation accuracy estimation for each window size (number of SNPs on each side) can be submitted simultaneously to the cluster. At each window size, NPUTE estimates imputation accuracy by looping through all known SNPs one by one, thus making it a time consuming process. A mismatch accumulator array (MAA) is adopted in NPUTE to improve performance, but parallelization can further reduce the computation time from days to hours.

**Fig. 4.**
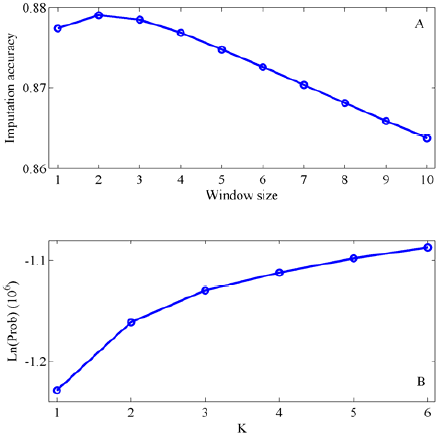
(A) Imputation accuracy at different window size; (B) Likelihoods of grouping data into K clusters.

Fig. 4A shows that the optimal window size is 2, which suggests that 2 SNPs on each side of the missing SNP offer the best information for the imputation. After this, the missing SNPs can now be imputed with the optimal window size determined in the estimation step. It is worthwhile to point out that the small window size of 2 is caused by highly linked SNPs in our processed data set (we choose to not filter SNPs by linkage disequilibrium or LD to keep more markers). In population genetics, LD is the non-random association of alleles at two or more loci [28]. Here, we choose to demonstrate NPUTE since it is also adopted by the original publication for processing sorghum data [23]. Certainly the iPlant CI is open for supporting additional imputation algorithms or utilizing pedigree information if available. After filling missing data (with NPUTE), filtering for minor allele frequency and conversion (using a tool named NumericalTransform, which is built using both TASSEL and PLINK [29]), the genotype data is ready for estimating population structure, or performing mixed model analysis using TASSEL, EMMAX, and MLMM.

We integrated STRUCTURE and parallelized it for simultaneously testing various assumptions of the number of populations (or clusters). Since STRUCTURE adopted a Markov Chain Monte Carlo (MCMC) Bayesian approach, we also allow the user to repeat each assumption several times simultaneously for checking whether the MCMC iteration numbers are sufficient for convergence. As an example, the likelihood (averaged across three repeats) value is plotted against each assumption of cluster numbers in Fig. 4B. For the SAP data, our integrated version of STRUCTURE allows quick estimation of population structure in hours instead of weeks. We provide a tool, named STRUCTURE4TASSEL, to format the output of STRUCTURE appropriately as input to TASSEL for GLM and MLM analysis.

It is tricky to interpret the optimal number of clusters from the likelihood values estimated by STRUCTURE. The general rule is to choose the one with the highest likelihood but not over-dividing the data into too many clusters. For re-analysis of SAP data, we assume the number of clusters is 3. Fig. 4B shows that the likelihood values are no longer increasing rapidly after assumption of 3 clusters (e.g. comparing the increases from 3 to 4 with 1 to 2 or 2 to 3). Furthermore, the outputs of STRUCTURE under different assumptions are kept so that the user can choose to further test each assumption with downstream tools, e.g. TASSEL.

For the SAP data, 310 accessions remained after merging genotype data with trait data (height). The output p values at each loci (x axis) from TASSEL (Fig. 5A-C), EMMAX (Fig. 5D) and MLMM (Fig. 5E), all after Bonferroni correction, are shown in Fig. 5. The dashed line on the right highlights the location of a dwarf gene, Dw1/SbHT9.1 on chromosome 9. Fig. 5E shows one stepwise regression results (MLMM model) after taking the most significant SNP (indicated by the dashed line on the right) as the co-factor to the mixed model. MLMM is designed to account for loci of larger effect, and interestingly adding the cofactor indeed highlights another SNP (55424715) that is not significant in a single point mixed model analysis (with either TASSEL or EMMAX). The SNP also falls on the coding part of a predicted transcript FGENESH00000020814 [31]. The nonsynonymous mutation (C/T) on this and adjacent loci causes the modest threonine (ACC) to isoleucine (ATC or ATT) conversion.

**Fig. 5.**
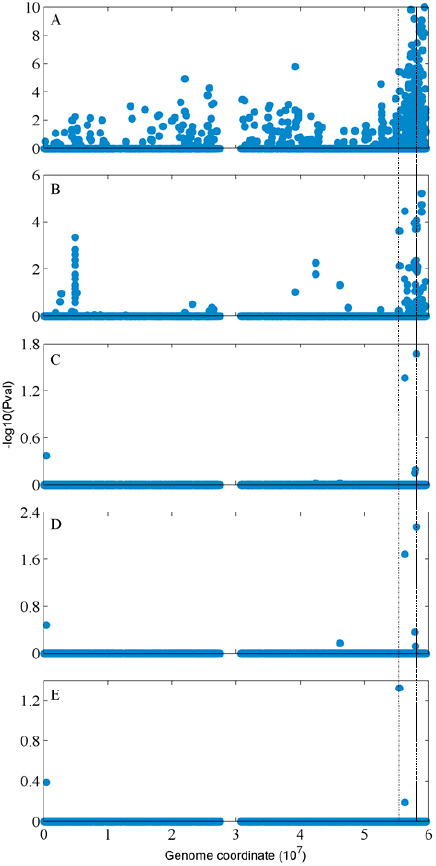
Output of various models with the dashed line highlighting the location of dw1/SbHT9.1 gene (right) and a predicted gene (left). (A) marker ∼ trait (TASSEL); (B) marker ∼ trait + population structure (TASSEL); (C) marker ∼ trait + kinship (TASSEL); (D) marker ∼ trait + kinship (EMMAX); (E) marker ∼ trait + kinship + cofactor (MLMM).

Further experimental verifications of all genes nearby might be necessary for confirming the above novel observation via MLMM. However, with this use case, we aim to demonstrate how a flexible GWA workflow is constructed on top of the iPlant platform. For this purpose, we have extended the original analysis by incorporating more popular (including STRUCTURE, EMMAX) or novel (MLMM) applications into the automated workflow. For format conversion needed among these applications, we choose to rely on existing implementations in well maintained applications, e.g. TASSEL and PLINK for sustainability.

In summary, the iPlant platform adds value to individual software applications by enabling the automation of GWA analysis, including the variant calling procedure (e.g. via GBS workflow), resolving accession names (via TNRS4GWAS), gaining more computing power to reduce computation time (e.g. via our parallelization of both NPUTE and STRUCTURE), and most importantly, simultaneously testing various assumptions and association models (e.g. TASSEL, EMMAX, MLMM) without worrying about format conversions. Another key advantage of the iPlant platform is that it is open for end users to integrate additional applications, modify existing applications, and build customized workflows.

### B. Nested Association Mapping (NAM)

Contrasting the SAP use case, where we assembled a collection of widely used applications for complex association analysis, in this NAM use case we aim to demonstrate the advantages of storing re-sequenced genomes to solve large-scale computation problems using the iPlant platform. The association analysis in this use case is limited to the GLM model with the bootstrapping method [30] implemented in TASSEL.

NAM was created as a method of combining the advantages and eliminating the disadvantages of two traditional methods for QTL (or linkage analysis) and association mapping. The maize NAM population was constructed from twenty-five diverse corn lines chosen as founder lines that encompass the remarkable diversity of maize and preserve the historic linkage disequilibrium. Each line was crossed to the B73 maize inbred to create the F1 population. The F1 plants were then self-fertilized for 6 generations to create a total of 200 homozygous RILs, for a total of 5000 RILs. Each RIL was originally genotyped with the same 1106 molecular markers [26], and extended to 7000 markers for 6000 RILs in this use case. The 25 founder lines were sequenced using NGS technology, and 1.6 million variable regions were initially discovered [31].

We have built an application, NAMGWAS, for GLM association of NAM SNPs with trait data uploaded by end users. This application is built with the SNP projection procedure, GLM modeling, and the bootstrapping method implemented in TASSEL. For this application, we hosted the SNPs for B73 and 25 founder lines, a NAM map file, and the 7000 markers for each NAM line. The NAM map file lists NAM markers with both genetic and physical positions, which are used to project the founder genotypes to the RILs using a nearest neighbor approach with window size of 1. The projection was done at runtime for each RIL at each locus for association with the trait of interest; then the projected SNP was discarded without storage. This turns out to be more efficient than storing all the SNPs for all RILs beforehand.

For the NAM genotype data set, it is not necessary to estimate the population structure since it is “known”: 25 populations corresponding to 25 founder lines. Therefore, the population information is integrated directly into the GLM model for reducing false positives. To quantify how significant a detected association is, the TASSEL implementation adopted a bootstrapping strategy, which turns out to be rather time consuming. Thus, we enhanced the NAMGWAS application for allowing a user to split bootstrapping iterations into multiple jobs at runtime for faster computation. Future optimization of the bootstrapping code in TASSEL will be critical for improving performance.

Using NAMGWAS, we were able to complete a large-scale NAM association analysis with 115 traits using over 1000 cluster nodes in a few weeks, rather than 2.5 years on a typical 8-core desktop. The data are unpublished, so results are not presented here. Going forward, the NAM strategy for storing genotypes might be adopted for other species other than maize, and they could be easily hosted by the iPlant platform given their much smaller size.

## VI. DISCUSSION

Association mapping with NAM data is straight forward given that their population structure is controlled by design. On the contrary, population structure within the SAP data needs to be estimated and corrected to effectively eliminate false positives. Looking at TASSEL outputs only, Fig. 5 shows that the false positive rate is significantly reduced after correcting for structure, and further reduced after correcting for kinship (finer scale of population structure). However, the tradeoff is that such corrections increase the chance of false negatives. For example, in Fig. 5, the larger value shown on the y axis indicates more significant association. Again, using the TASSEL outputs as an example, the most significant association decreases from 9 to 5 in value after correction for structure effect, and further reduces to 1.6 after kinship correction. This has created another problem that the true association might not be detected if the threshold is not set appropriately. In the worst case scenario, when the trait of interest completely overlaps with the population structure, the true association will be erased by correction of structure effects. Therefore, it is always better to test multiple models and assumptions. This is also one of the rationales why we have constructed an automated workflow for simultaneously testing different models. It has also been suggested that the combination of traditional linkage analysis [32] and further haplotype analysis [33] can increase the power of GWA mapping to distinguish true from false associations.

On the other hand, the stepwise regression adopted by the MLMM approach is simply a greedy forward-backward search strategy. The ideal solution will be evaluating all combinations of SNPs, which is usually impossible for the dense SNPs available today since the number of combinations increases with the number of SNPs exponentially. An alternative solution or improvement will be reducing the search space by restricting the combination tests to SNPs close to (or linked to) genes involved in the related pathways. For new genomes, this will be related to iPlant’s genome assembly efforts to construct reference genomes, and annotation efforts to identify true genes. We have also developed in-house applications for *de novo* gene prediction based on the modified wavelet transform [34], and examination of exon sizes and their evolution [35, 36] that can improve annotation accuracy.

Again, iPlant’s mission is beyond GWA. In this work, we aim to provide a “GWAS view” of iPlant CI for handling complicated analysis workflows. In fact, the iPlant CI development in the first five-year grant period has been focused on developing an open system for application integration, leveraging high performance computation, and large scale un-relational data management. As demonstrated here, these goals have been extremely successfully accomplished. However, much more sophisticated data modeling is key to extend or combine GWA to or with iPlant’s broad efforts in NGS analysis, pathway integration, assembly, and annotation. Therefore, one of iPlant’s main goals for the second five-year grant period is to extend the data store to provide published data sets bringing data closer to computation for user. This will include supporting storage and accessing of variations (SNPs, indels, structural variations, etc.) in VCF (Variant Call Format) [37]. The extended data store will also facilitate data transfer between iPlant and public data repository (e.g. pathway database), provide quality control (QC), support image based phenotyping [21, 38], and enhance visualization.

On the technical side, the GWA related applications we integrated into iPlant CI for executing on high performance clusters are developed with a wide variety of programming languages, including C++, Java, R, Perl, Matlab, python, and even shell scripts. This reduces the efforts in rewriting applications for integration into an application package, and demonstrates that the iPlant platform is flexible for end users to plug in their own or favorite applications to utilize the high performance computing clusters. However, the challenges for building an open system are usually much bigger than building a closed system. Such CI challenges, in some sense, have also been complicating the scientific developments in the last few years. We believe that the further development of the iPlant platform will make it continuously maturing in both stability and functionality.

## ACKNOWLEDGMENT

This work is supported by the National Science Foundation Plant Cyber-infrastructure Program (#DBI-0735191 and #DBI-) and USDA-ARS. The authors would like to thank the two anonymous reviewers for their helpful comments. L. W. would like to thank Drs. Joshua Stein and Stephen Goff for critical reading of the paper, Drs. Peter Bradbury and Ed Buckler for the early collaboration on integrating TASSEL, Terry Casstevens for collaboratively developing the command line plugins for the TASSEL suites (basis for integrating TASSEL into iPlant CI), Dr. Qi Sun for helping on integrating GBS workflow and analyzing sorghum data, Dr. Kang for helping on EMMAX integration, Dr. Geoffrey Morris for assisting with extracting the sorghum data, Dr. Ivan Baxter and Greg Ziegler for the NAM bootstrapping collaboration, Drs. Rion Dooley, Matthew Vaughn, and Paul Navratil for assistances on deploying apps on TACC Lonestar and Longhorn clusters.

